# IVISc-L: A quick and simple *in vivo* assay to study the regulation of gene expression

**DOI:** 10.1101/2024.12.16.628807

**Authors:** Saubhik Som, Gopalapura J Vishalakshi, Lekha E Manjunath, Debraj Manna, Kirtana Vasu, Anumeha Singh, Sandeep M Eswarappa

## Abstract

Several methods are available to study the regulation of gene expression at cellular and molecular levels. Adaptation of these methods *in vivo* is cumbersome and often requires animal sacrifice. Here, we report an assay (IVISc-L, *In Vivo* Imaging of Subcutaneous Luminescence) to study gene regulation *in vivo*. This assay involves subcutaneous injection of a plasmid DNA encoding firefly luciferase, whose expression is under the regulatory mechanism to be investigated. We could infer its regulated expression by detecting the subcutaneous luminescence using an *in vivo* imaging system. Using this assay, we have demonstrated the regulation of gene expression mediated by a promoter, micro-RNAs, stop codon readthrough, and rare codons. This minimally invasive assay does not require animal sacrifice or any tissue extraction. The entire assay can be completed within 24 hours. Therefore, this assay will be useful in investigating the mechanisms of gene expression regulation, and screening molecules that can alter gene expression *in vivo*.

## INTRODUCTION

Temporal and spatial regulation of gene expression is vital for cells to perform their functions. They govern many cellular processes including development, metabolism, homeostasis, immune functions among many others [1]. Aberrations in gene expression can lead to pathological conditions such as cancer [2]. Gene expression is regulated at multiple levels from transcription to protein degradation. Several molecular methods have been developed to study the regulation of gene expression at the global level as well as the individual gene level. Incorporation of radioactive or fluorescent amino acids (e.g., [^35^S]Methionine) [3], RiboPuromycylation [4], click chemistry-based assays [5], RNA sequencing [6], ribosome profiling [7], and quantitative mass spectrometry [8] are extensively used to study gene expression changes at various levels. To investigate the regulation of expression of a specific gene, fluorescence and luminescence-based reporter assays, quantitative RT-PCR, and western blotting are used.

Many of these techniques can also be used *in vivo* to study regulation of gene expression in animal models. However, they require extraction of tissues by sacrificing the animal, which makes such assays lengthy, expensive, and cumbersome. Therefore, there is a need for quicker, easier, and inexpensive methods. Here, we report a simple and relatively quick *in vivo* method to study the regulation of gene expression. This involves subcutaneous injection of naked DNA in mice.

The very first report of expression of exogenous proteins from injection of naked DNA *in vivo* came in 1990. Injection of naked plasmids encoding chloramphenicol acetyltransferase, luciferase, and beta-galactosidase into mouse skeletal muscles resulted in expression of these proteins, which were detected by activity assays in muscle tissue samples [9]. This study played a pivotal role in the emergence of DNA vaccines. Here, we have employed subcutaneous naked DNA injection to express luciferase reporter in mice whose activity could be detected using a sensitive *in vivo* imaging system without sacrificing the mice. We show that this relatively quick and minimally invasive method, termed IVISc-L (*In <u>V</u>ivo* Imaging of <u>S</u>ub<u>c</u>utaneous <u>L</u>uminescence), can be used to investigate different modes of regulation of gene expression.

## RESULTS AND DISCUSSION

### Naked DNA injection results in signal detectable by *in vivo* optical imaging system

We tested whether naked DNA injection in mice can generate exogenous proteins that can be detected without harming the animal. For this, we used a plasmid that encodes firefly luciferase. This enzyme generates luminescence when its substrate luciferin is provided. Luminescence can be detected using an optical imaging system. Instead of using just firefly luciferase, we tagged its coding sequence (CDS) to partial CDS of *MTCH2* that encodes a mitochondrial carrier protein. This is because, the coding sequence (CDS) of firefly luciferase is usually tagged to a test sequence in reporter assays to study gene regulation, (Fig 1A). We injected the naked plasmid (10 μg) encoding this tagged firefly luciferase in mice via multiple routes – intraperitoneal, intravenous, subcutaneous at the flanks, and subcutaneous in the tail. After 24 hours, these mice were injected with D-luciferin and imaged using IVIS spectrum *in vivo* imaging system. We were able to detect a robust luminescence signal in mice injected subcutaneously in the tail (Fig 1A). Next, to know the optimal dose required to obtain detectable signal, we injected the reporter construct at multiple doses and performed the same assay. A dose of 10 μg was sufficient to obtain a robust luminescence signal by the imager (Fig 1B). The luminescence signal was detectable as early as 10 hours and stayed up to 72 hours after the injection. However the signal was much reduced by 72 hours (Fig 1C). We next tested whether this assay, termed *In Vivo* Imaging of Subcutaneous Luminescence (IVISc-L), can be used to study different modes of regulation of gene expression. To get robust and consistent signal, we used reporter plasmids at a dose of 20 μg/mouse, and the *in vivo* imaging was done 24 hours after the injection in subsequent experiments.

**Figure 1.**
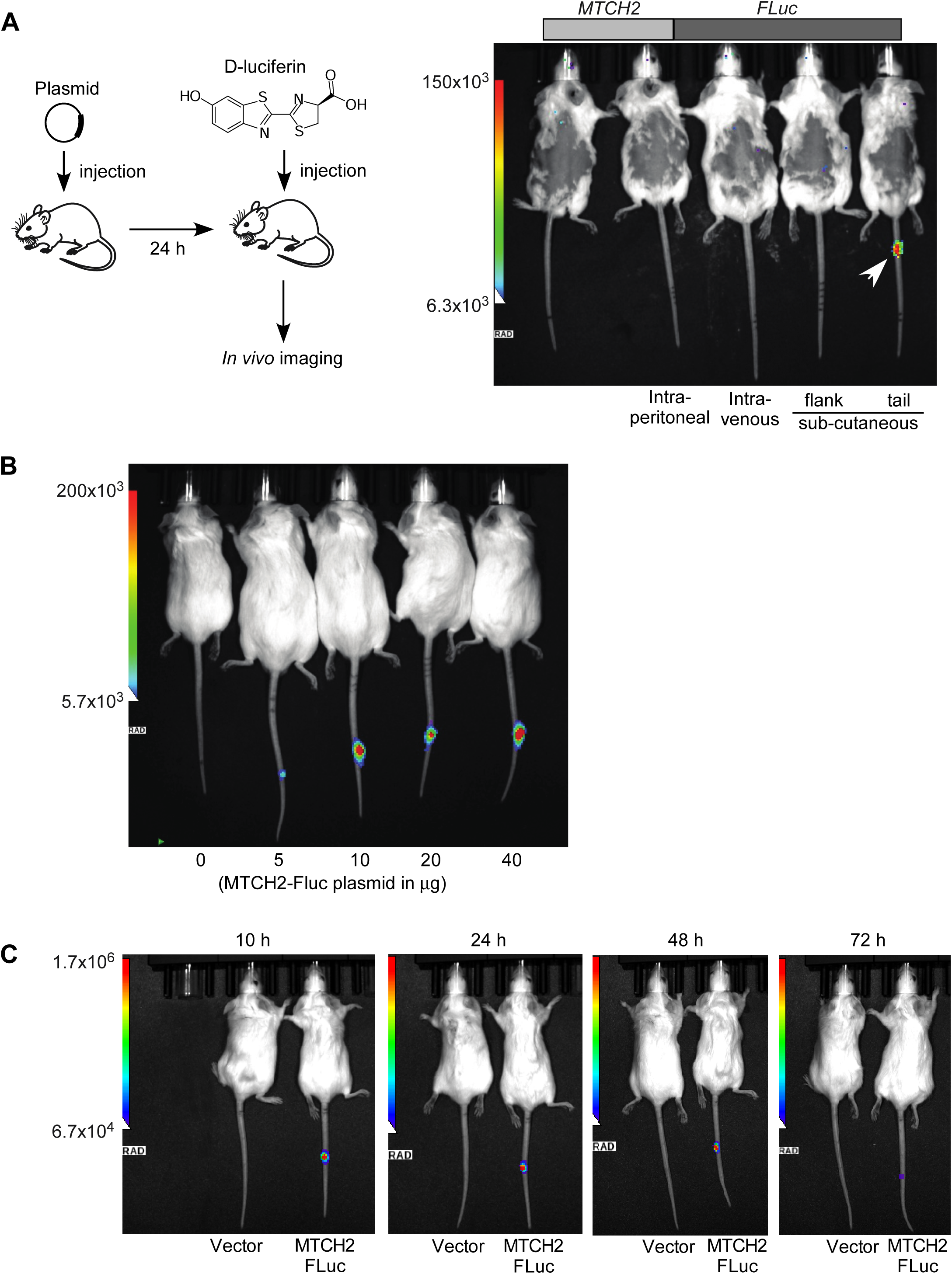
Detection of luminescence resulting from naked reporter DNA injection. (A) A plasmid expressing firefly luciferase was injected via indicated routes (dose, 10 μg/mouse). After 24 h, mice were injected with D-luciferin (150 mg/kg, intraperitoneally) and anesthetized for detecting the luminescence signal using IVIS spectrum *in vivo* imaging system. The exposure time was 60 sec. Intensity of the signal is shown (6.3×10^3^ to 150×10^3^ photons/sec). (B) Intensity of the luminescence signal (5.7×10^3^ to 200×10^3^ photons/sec) in mice injected with the plasmid reporter expressing firefly luciferase via subcutaneous route in tail. Dose, 0 to 40 μg/mouse. (C) Intensity of the luminescence signal in mice 10, 24, 48, and 72 h and after the subcutaneous injection of the reporter plasmid in tail. Dose, 20 μg/mouse

### Detection of regulation of gene expression at the level of transcription

Regulation of gene expression at the level of transcription is important in all cellular processes and functions [10]. We first investigated if IVISc-L can detect the differences in gene expression at the level of transcription. For this, we cloned the promoter and enhancer of cytomegalovirus (CMV) upstream of firefly luciferase (FLuc) in pGL3 promoter less vector (Fig 2A). When transfected in HeLa cells, the construct with promoter showed robust firefly luciferase activity, while the control construct without the promoter showed only background activity, as expected (Fig 2B). After confirming differential expression in cells, we performed IVISc-L assay in mice. Mice injected with construct having CMV promoter showed strong luminescence signal, and those injected with the control construct showed only background signal (Fig 2C). These results show that IVISc-L assay can be used to study the regulation of gene expression at the level of transcription.

**Figure 2.**
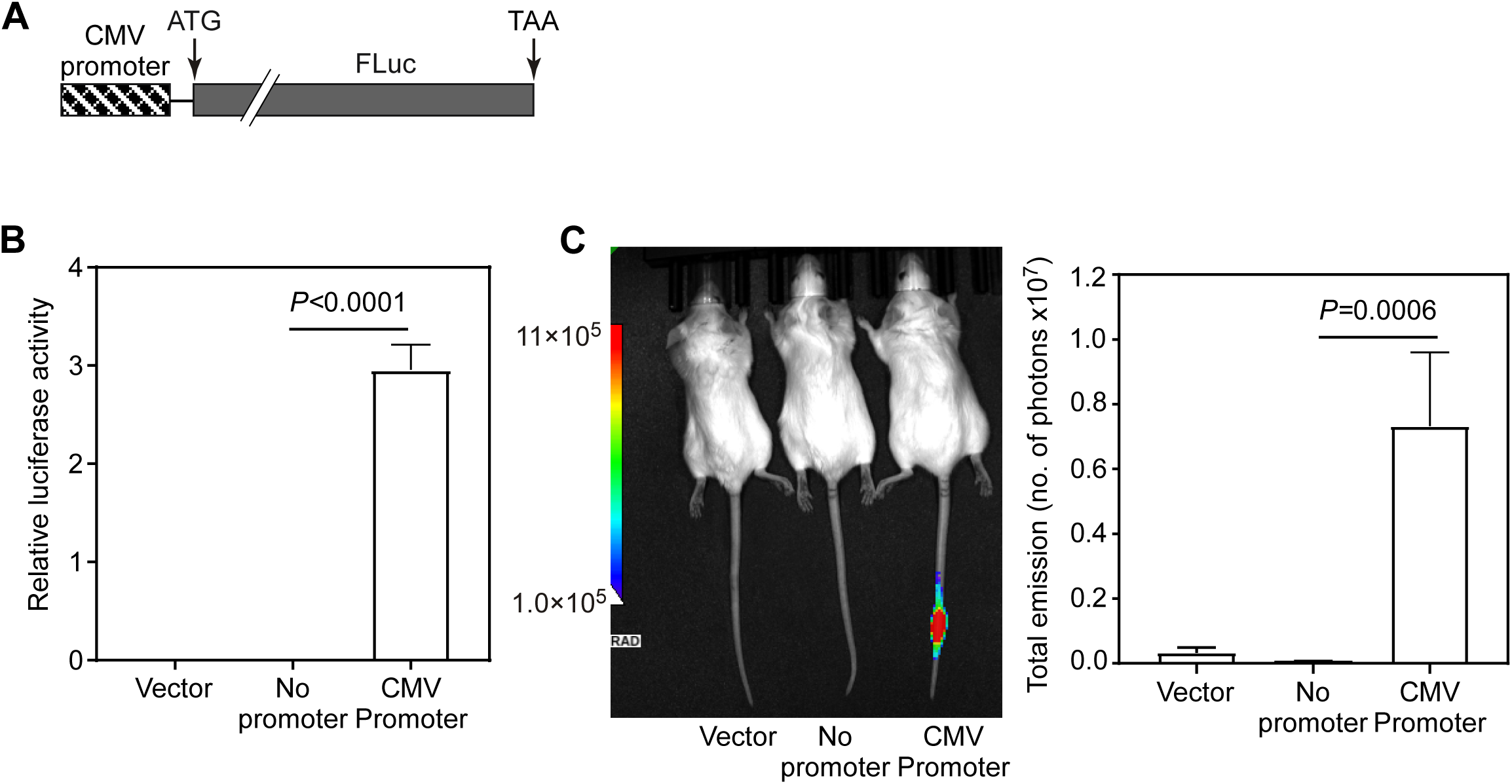
Detection of regulation of gene expression at the level of transcription. (A) Schematic of the luciferase construct used in the IVISc-L assay. The CMV promoter was included upstream of the CDS of firefly luciferase in pGL3 vector backbone. (B) Luciferase assay in HeLa cells transfected with indicated constructs. Firefly luciferase activity relative to the activity of co-transfected *Renilla* luciferase is plotted. Mean ± SE, N=3. *P* value, two-tailed student’s t-test. (C) IVISc-L assay: Intensity of the luminescence signal (1×10^5^ to 11×10^5^ photons/sec) in mice injected with the plasmid reporter shown in (A) via subcutaneous route in tail. Dose: 20 µg/mouse. Mean ± SE, N=7 from two independent experiments.

### Detection of microRNA-mediated regulation of gene expression

Next, we tested the ability of IVISc-L assay to detect the regulation of gene expression at post-transcriptional level. MicroRNAs (miRNAs) are small noncoding RNAs that target messenger RNAs. In mammals they usually target 3′ untranslated regions (UTRs) of mRNAs and bring down their translation and/or induce their degradation [11]. We employed this pathway to test the ability of our *in vivo* method to detect gene regulation at post-transcriptional level. The CDS of firefly luciferase was tagged with the proximal 3′UTR (897 nucleotides) of *PDCD4*, which is a known target of multiple microRNAs - miR-21, miR-182, miR-320a, miR-183, and miR-499-5p [12–18]. As a control, we used a sequence of the same length (897 nucleotides), which does not have any known or predicted microRNA targets, in place of the 3′UTR of *PDCD4* (Fig 3A).

**Figure 3.**
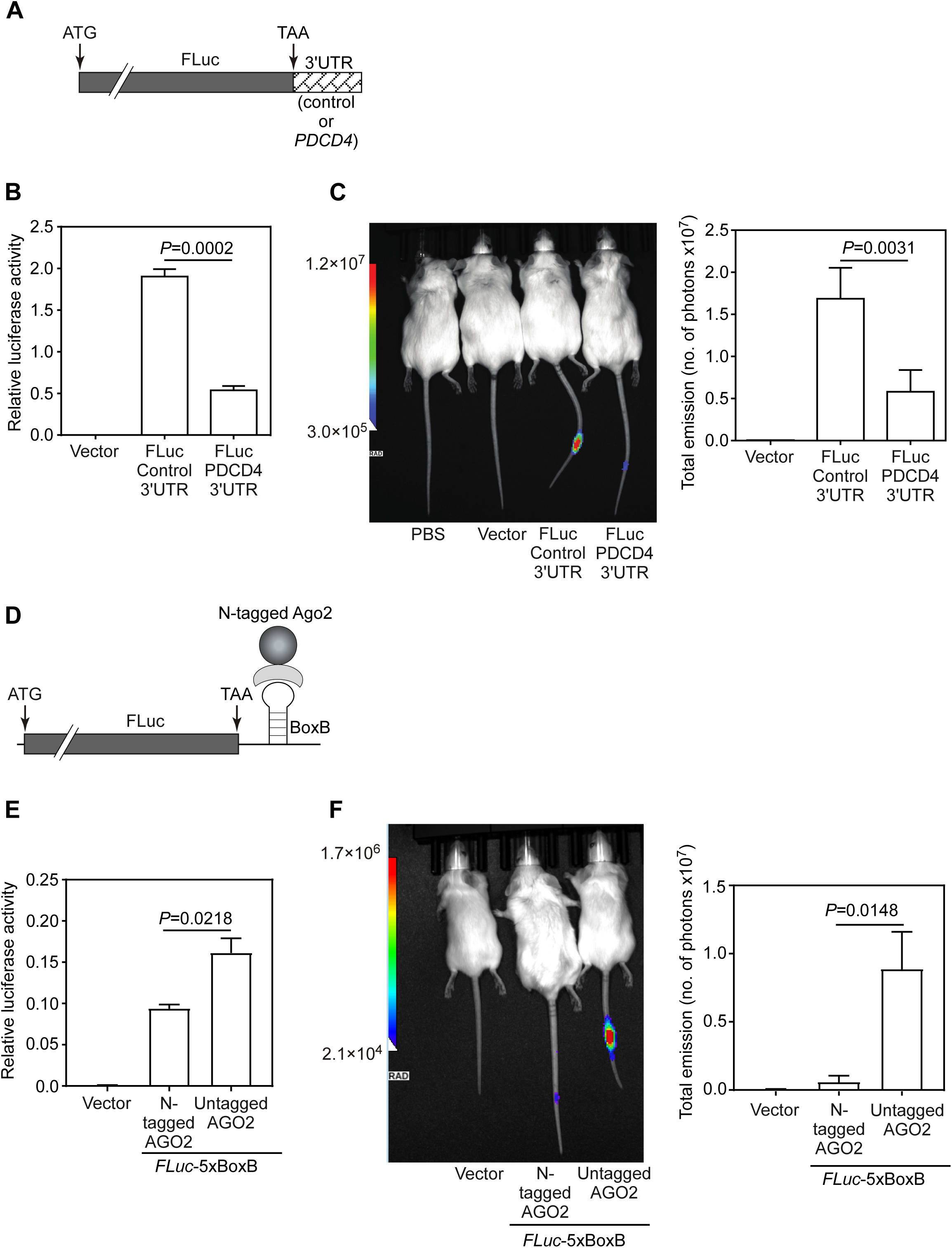
Detection of microRNA-mediated regulation of gene expression using IVISc-L assay. (A) Schematic of the luciferase construct used in the IVISc-L assay. The 3′UTR of *PDCD4* was appended to the CDS of firefly luciferase. (B) Luciferase assay in HeLa cells transfected with indicated constructs. Firefly luciferase activity relative to the activity of co-transfected *Renilla* luciferase is plotted. Mean ± SE, N=3. *P* value, two-tailed student’s t-test. (C) IVISc-L assay: Intensity of the luminescence signal (3×10^5^ to 120×10^5^ photons/sec) in mice injected with the plasmid reporter shown in (A) via subcutaneous route in tail. Dose, 20 μg/mouse. Mean ± SE, N=19 for control and N=17 for test. Mice were pooled from four independent experiments. *P* value, Mann-Whitney U-test. (D) Schematic of the BoxB-N-peptide tethering assay used. Five BoxB motifs, which form stem-loop structures, were appended at the 3’UTR of the CDS of firefly luciferase. Ago2 tagged with N-peptide from bacteriophage λ will interact with BoxB elements, recruit RNA induced silencing complex and bring down the expression of firefly luciferase. (E) Luciferase assay in HeLa cells transfected with indicated constructs. Firefly luciferase activity relative to the activity of co-transfected Renilla luciferase is plotted. Mean ± SE, N=3. (F) IVISc-L assay: Intensity of the luminescence signal (2.1×10^4^ to 170×10^4^ photons/sec) in mice injected with the indicated constructs via subcutaneous route in tail. Dose, *Fluc-*5×BoxB: 10 µg/mouse; N-tagged AGO2/Untagged AGO2: 20 µg/mouse. Mean ± SE, N=7 for both from two independent experiments.

To make sure that these constructs show differential expression, they were first transfected in HeLa cells. The firefly luminescence was measured relative to the co-transfected *Renilla* luciferase. As expected, we observed reduced expression of firefly luciferase in the constructs with the 3′UTR of *PDCD4* compared to the control (Fig 3B). We then injected these constructs subcutaneously in the tails of mice and the luminescence signal was quantified using the *in vivo* imaging system. We observed more than 50% reduction in the signal in mice injected with the construct having the 3′UTR of *PDCD4* compared to the control (Fig 3C).

BoxB-N-peptide tethering system is a very useful tool to investigate miRNA-mediated regulation of gene expression at post-transcriptional level [19]. This system is derived from bacteriophage λ. The stem-loop RNA element BoxB interacts with a 22-amino acid-long peptide called N-peptide, which is important for transcription anti-termination in these bacteriophages. We cloned five BoxB elements downstream of the CDS of firefly luciferase (Fig 3D). When this construct was expressed in HeLa cells along with N peptide-tagged Ago2, the expression of firefly luciferase was reduced when compared to cells expressing Ago2 without the N peptide tag (Fig 3E). This result showed that the assay is working as expected. We then performed IVISc-L assay using these constructs. Similar to HeLa cells, we observed reduced firefly luciferase activity in mice injected with five BoxB firefly luciferase and N peptide-tagged Ago2 construct compared to mice injected with Ago2 without the tag (Fig 3F). These results show that the IVISc-L assay can be used to investigate micro RNA-mediated post-transcriptional gene regulation.

### Detection of translational readthrough across stop codons

Our next goal was to test the regulation of gene expression at the level of translation using IVISc-L assay. For this, we chose to study stop codon readthrough. In certain mRNAs, translating ribosomes continue translation beyond stop codons, instead of terminating at that position. This process, termed as stop codon readthrough (SCR), generates C-terminally extended isoforms. Because of the unique C-terminus, the SCR isoforms can have different function, localization and stability compared to the canonical isoform [20]. Mammalian *MTCH2*, which encodes a mitochondrial carrier protein, exhibits SCR. This process is important to maintain normal mitochondrial membrane potential [21]. We investigated if the IVISc-L assay can be used to demonstrate SCR of *MTCH2*. The CDS of firefly luciferase was cloned downstream of and in frame with the partial CDS and the proximal 3′UTR (45 nucleotides) of *MTCH2* as described previously [21](Fig 4A). In this construct, luciferase is expressed *only* if there is translational readthrough across the stop codon of *MTCH2* as the start codon of luciferase is absent. Stop codon readthrough was first confirmed by transfecting this construct in HeLa cells. A construct without the proximal 3′UTR of *MTCH2*, which is the readthrough-driving element, was used as a negative control. We observed a clear and significant luminescence signal in the construct with the readthrough-inducing element (Fig 4B). To investigate this phenomenon *in vivo*, these constructs were injected subcutaneously in mouse tail as described above. Similar to the results in HeLa cells, we observed stop codon readthrough-induced luminescence in mice. This signal was not detected in mice injected with the construct that lacks the readthrough-inducing element (Fig 4C). These results show that IVISc-L assay can be used to study the regulation of gene expression by SCR.

**Figure 4.**
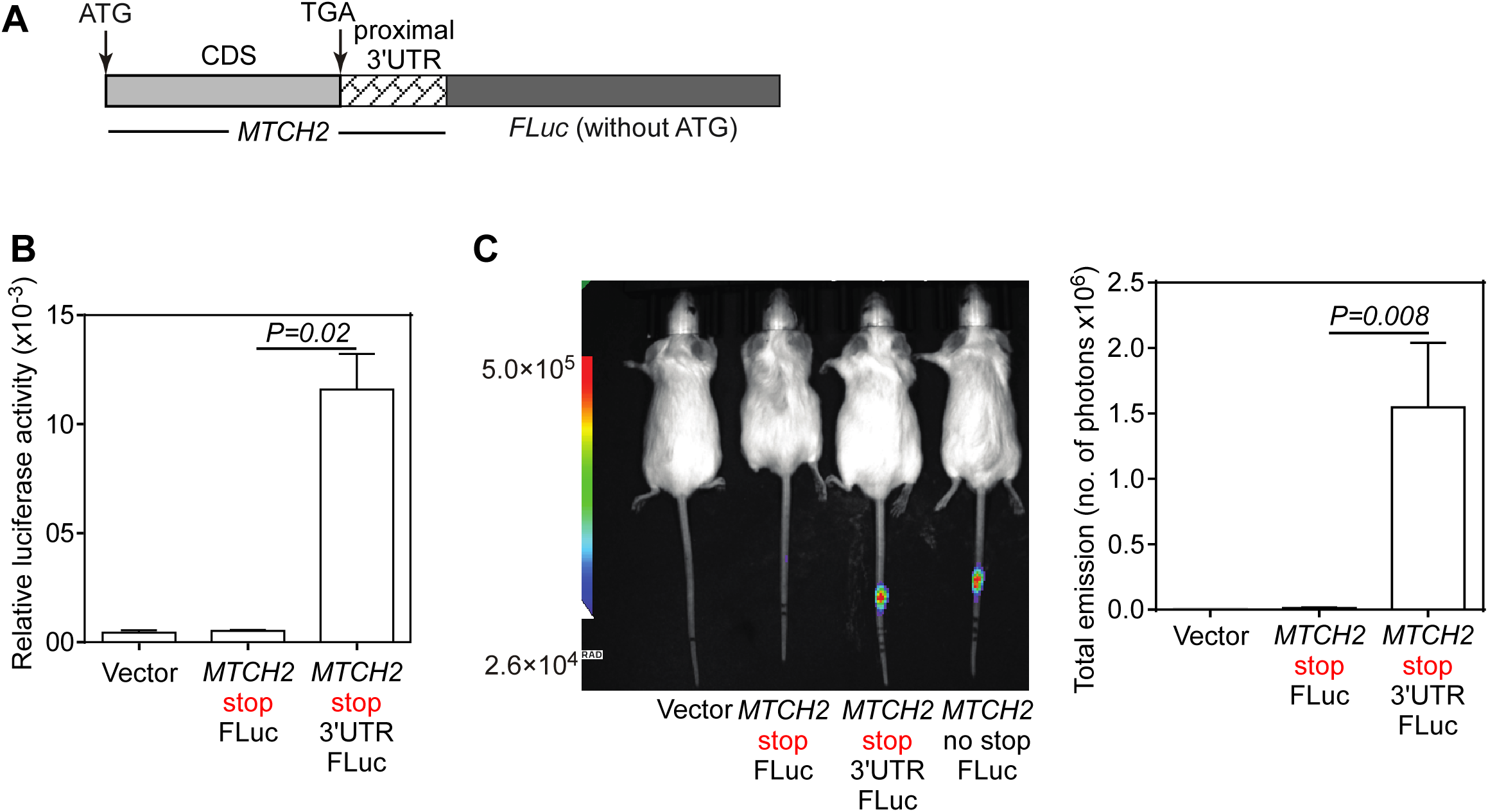
Demonstration of stop codon readthrough of *MTCH2* using IVISc-L assay. (A) Schematic of the luciferase construct used in the IVISc-L assay. The CDS of firefly luciferase was cloned downstream of the CDS and the proximal 3′UTR of *MTCH2* (45 nucleotides) such that firefly luciferase is expressed only if there is stop codon readthrough across the stop codon of *MTCH2*. (B) Luciferase assay in HeLa cells transfected with indicated constructs. Firefly luciferase activity relative to the activity of co-transfected *Renilla* luciferase is plotted. Mean ± SE, N=3. *P* value, two-tailed student’s t-test. (C) IVISc-L assay: Intensity of the luminescence signal (2.6×10^4^ to 50×10^4^ photons/sec) in mice injected with the plasmid reporter shown in (A) via subcutaneous route in tail. Dose, 20 μg/mouse. Mean ± SE, N=8 from four independent experiments. *P* value, Mann-Whitney U-test.

### Detection of rare codon mediated translational regulation

Many amino acids are encoded by more than one codon. The frequency of occurrence of these codons in the transcriptome of an organism is not uniform. Some of them occur more frequently and some of them rarely. Presence of rare codons slows down the rate of the translation elongation as the corresponding tRNAs are also less abundant. This slowing down of translation influences mRNA stability, protein folding, and even its function [22]. We investigated if the IVISc-L assay could detect translational regulation by codon usage.

We inserted 16 rare codons (CTT CAA ATA GTA CTC GCA ATT GTT CTC GCG GCT CAA ATT GTC CAA GTA) near the 5′ end of the CDS of firefly luciferase. Upstream to these rare codons, we included 8 optimal codons (CTG CAG ATC GTG CTG GCC ATC GTG) to make sure that the initiation is not directly affected by these rare codons. As a control, we replaced the 16 rare codons by 16 synonymous optimal codons (CTG CAG ATC GTG CTG GCC ATC GTG CTG GCC GCC CAG ATC GTG CAG GTG) (Fig 5A). We first transfected these constructs in HeLa cells and measured the firefly luciferase activity relative to the activity of the co-transfected *Renilla* luciferase. Expectedly, we observed the rare codon-induced reduction in the expression of firefly luciferase (Fig 5B). We then injected these constructs subcutaneously in the mouse tail and measured the luminescence using the *in vivo* imager. In agreement with the results of the cell-based assay, we observed reduction in the activity of firefly luciferase containing rare codons (Fig 5C). This shows that IVISc-L assay can be used to detect the changes in gene expression caused due to rare codons.

**Figure 5.**
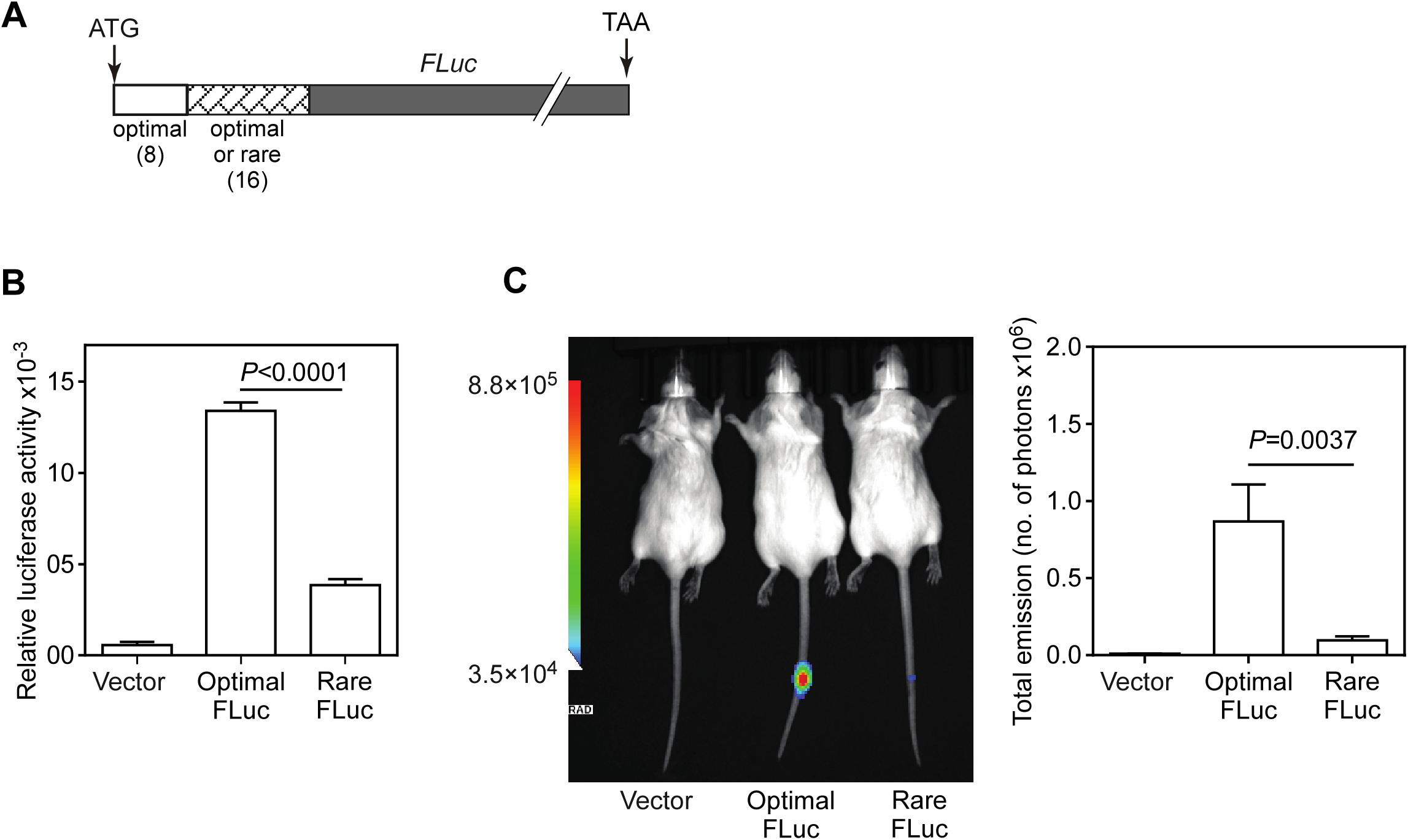
Demonstration of rare codon-mediated regulation of translation. (A) Schematic of the luciferase construct used in the IVISc-L assay. Sixteen rare codons or their corresponding optimal codons were appended to the 5′ end of the CDS of firefly luciferase. (B) Luciferase assay in HeLa cells transfected with indicated constructs. Firefly luciferase activity relative to the activity of co-transfected *Renilla* luciferase is plotted. Mean ± SE, N=3. *P* value, two-tailed student’s t-test. (C) IVISc-L assay: Intensity of the luminescence signal (3.5×10^4^ to 88×10^4^ photons/sec) in mice injected with the plasmid reporter shown in (A) via subcutaneous route in tail. Dose, 20 μg/mouse. Mean ± SE, N=10 for optimal codon condition and N=11 for rare codon condition. Mice were pooled from two independent experiments. *P* value, Mann-Whitney U-test.

*In vivo* studies in animal models are required to confirm the cellular processes observed *in vitro* in cell lines. Our experiments demonstrate the utility of IVISc-L assay in the study of gene expression *in vivo*. In this assay, we have utilized the previous observation that naked DNA injection can generate proteins [9]. Using the highly sensitive *in vivo* optical imaging system, we were able to detect the luminescence emanating from the subcutaneous region of mouse tail injected with naked reporter DNA. Notably, the signal was detectable only after subcutaneous injection in the tail region. Other locations and routes did not provide detectable signal. It is possible that the cells in the tail are more receptive for the naked DNA and/or the luminescence signal can effectively penetrate the skin barrier in the tail. These possibilities remain to be investigated. Our observations also provide support for the possibility of delivering nucleic acid-based drugs via subcutaneous route for dermatological conditions [23].

Our experiments demonstrate that IVISc-L assay can be used to investigate the regulation of gene expression at transcriptional, post-transcriptional and translational levels. As it involves just a subcutaneous prick and inhalation anaesthesia, this technique is minimally invasive in nature. Furthermore, the entire process can be finished within 24 hours, which is comparable to assays performed in cell culture. As the signal disappears in a few days after injection, the mice can be reused. These advantages make it a very convenient and inexpensive assay to study gene expression mechanisms *in vivo*. The assay can also be scaled up and used to screen for drugs that can modulate gene expression *in vivo*.

## MATERIALS AND METHODS

### Plasmid construction

To optimize the route, dose, and time, firefly luciferase tagged to MTCH2 at the N-terminus was used. Partial coding sequence of *MTCH2* (732 nucleotides at its 3′ end without its stop codon) was cloned (between *Hind*III and *Bam*HI sites) upstream of and in-frame with the CDS of firefly luciferase (FLuc) without its start codon [21]. The construct used to test stop codon readthrough of *MTCH2* has been described previously [21]. To investigate the microRNA-mediated regulation, 5′ 897 nucleotides of the 3′UTR of *PDCD4* was cloned downstream of the CDS of firefly luciferase between *BamH*I and *Xho*I sites. As a control, the remaining 3′UTR (897 nucleotides) of *PDCD4* was cloned between same enzyme sites. In another construct used to test micro-RNA mediated regulation, reversed sequence of 5x-BoxB from pCIneo-RL-5BoxB [19] was cloned downstream of firefly luciferase CDS between *Xho*I and *Xba*I sites in pcDNA3.1 backbone. pIRESneo-FLAG/HA-Ago2 (without N-peptide tag) and pCIneo-N-HA-Ago2 (with N-peptide tag) are described in previous studies [19, 24]. To observe the effect of rare codons, 16 rare codons (or synonymous optimal codons as control) were cloned upstream of firefly luciferase CDS between *Hind*III and *BamH*I sites. For promoter activity, CMV promoter from pcDNA3.1myc-His B was cloned between *Xho*I and *Hind*III sites upstream of the CDS of firefly luciferase in pGL3 vector. Sequences of all the constructs used in this study are provided in Supplementary Information.

### Mice experiments

Mice experiments were conducted after the approval from Institutional Animal Ethics Committee (CAF/ETHICS/066/2024). The reporter plasmid constructs were dissolved in 30 μl of phosphate buffered saline and injected via intravenous route (tail vein) or intraperitoneal route or subcutaneous route (flank or tail) using a syringe (U40 from BD) with ultra fine needle (31-gauge) under aseptic conditions. After indicated time points, mice were injected with 150 mg/kg of D-luciferin (GOLDBIO) intraperitoneally using a 1-ml syringe and 26-gauge needle (Dispo Van). Ten minutes after the injection, mice were anaesthetized using 2.5% isoflurane in Rodent Anaesthesia System RAS-4 (PerkinElmer) and the luminescence signal was detected using IVIS spectrum *in vivo* imaging system (PerkinElmer) with an exposure time of 60 seconds. Flanks of mice were shaved to increase the chances of detection of luminescence from the body of the mice in case of injection via intravenous and intraperitoneal routes. The images were analysed using Aura 3.2 to quantify the emission (photons/sec) from the region of interest.

### Luciferase assays in HeLa cells

HeLa cells were maintained in Dulbecco’s modified Eagle’s medium (HiMedia) containing 10% fetal bovine serum (Thermo Fisher Scientific), 100 units/ml penicillin, and 100 μg/ml streptomycin (HiMedia) at 37°C in a humidified atmosphere containing 5% CO2. Cells were authenticated by STR profiling. They were checked for mycoplasma contamination twice a year. Cells at around 70% to 80% confluency in a 24-well plate were transfected with 500 ng/well of plasmid constructs expressing firefly luciferase using 4 ng/µl of polyethyleneimine (Polysciences, inc.). A plasmid expressing *Renilla* luciferase was co-transfected (50 ng/well), which served as transfection control. Cells were lysed with passive lysis buffer (Promega) 24 hours post-transfection, and luciferase activity was measured using Dual-Luciferase Reporter Assay System (Promega) in the GloMax Explorer System (Promega).

### Statistics

In assays with HeLa cells, two-tailed student’s t-test was performed. Welch’s correction was applied wherever equal variance was not observed. Mann-Whitney U-test was performed for IVISc-L assays. Outliers with background-subtracted signal beyond mean ± 2 SD were excluded from the analyses.

## ACKNOWLEDGEMENTS

We thank the Central Animal Facility the of Indian Institute of Science, and its animal imaging facility. This work was supported by the financial support from the Swarnajayanti Fellowship (DST/SJF/LSA-04/2019-20) given by the Department of Science and Technology (DST) and Science and Engineering Research Board, STARS grant from the Ministry of Education (STARS/APR2019/BS/328/FS), DBT’s Genome Engineering Technology program (BT/PR38405/GET/119/309/2020), EMBO Global Investigator Network, DST Funds for Improvement of S&T infrastructure, and the Institute of Eminence funds given by the Ministry of Education to Indian Institute of Science. SS acknowledges Prime Minister Research Fellowship provided by Government of India and LEM acknowledges the Research Associateship received from the Indian Council for Medical Research (ID-2021-13048).

## Conflict of Interest

The authors declare that they have no conflict of interest

